# Biosynthesis of the proteins containing neurotoxin *β*-*N*-methylamino-L-alanine in marine diatoms

**DOI:** 10.1101/2023.12.21.572844

**Authors:** Xianyao Zheng, Aifeng Li, Jiangbing Qiu, Guowang Yan, Peng Zhao, Min Li, Fanping Meng

## Abstract

Neurotoxin *β*-*N*-methylamino-L-alanine (BMAA) has been deemed a pathogenic factor for human neurodegenerative diseases. It is an important issue to disclose the biosynthesis mechanism of BMAA in marine diatoms. In the present study, the iron (Fe) limitation (1/3 × Fe) was found to suppress the growth of diatoms but stimulate the production of BMAA-containing proteins, maximum 7.7 fold in *Thalassiosira minima*. Transcriptome analysis showed that energy metabolism, protein biosynthesis and carbon fixation functions were mainly affected by the Fe limitation in the diatom. Analysis of subcellular distribution of BMAA showed that BMAA-containing proteins were mainly detected in the endoplasmic reticulum and the Golgi apparatus. Combination results of the responses of the diatom to Fe deficiency and co-culture with cyanobacteria in our previous study, we speculate that cysteine embedded in peptide chains and methylamine produced by the diatom itself are possibly catalyzed by the cysteine synthase (cysK) to form the BMAA structure in situ. Spiked methylamine in culture media significantly stimulated the production of BMAA, and BMAA amounts were correlated with the expression of cysK gene in different diatoms. The reduced ubiquitination-mediated proteolysis and vesicle trafficking precision through the COPII system would aggravate the accumulation of BMAA-containing proteins in the diatom.

**Significance Statement:** With the detection of neurotoxin BMAA in diverse marine diatoms, the pathogenic risk of BMAA has been further concerned to human neurodegenerative diseases such as Alzheimer’s disease. Interestingly, BMAA-containing proteins are the dominant forms of this neurotoxin in diatoms. It is a keystone issue to disclose the biosynthesis mechanism of BMAA in marine diatoms. We found Fe-limitation could stimulate the production of BMAA-containing proteins in diatoms and explored its biosynthesis using transcriptomics in this study. Results suggested that cysteine embedded in peptides and methylamine in cytoplasm were catalyzed by the cysteine synthase (cysK) to form BMAA. This study hints that the biosynthesis of BMAA would be improved by the worldwide prevalence of iron deficiency in the coastal waters.

## Introduction

The neurotoxic *β-N*-methylamino-L-alanine (BMAA) was deemed an environmental causative agent for the neurodegenerative diseases such as amyotrophic lateral sclerosis/ parkinsonism dementia complex (ALS/PDC), and Alzheimer’s disease (Cox et al. 2003). In recent years, BMAA was detected in marine phytoplankton including diatoms (Réveillon et al. 2016; Wang et al. 2021) and dinoflagellates (Jiang and Ilag 2014; Lage et al. 2014). Bioaccumulation and transfer of BMAA to higher trophic levels are responsible for the human exposure to BMAA and thus its potential impact on human health (Nunes-Costa et al. 2020; Wang et al. 2021). It was alarming that the increased exposure of BMAA *via* food chains to humans can occur from marine diatoms (Esterhuizen-Londt and Pflugmacher 2019). However, the biosynthesis mechanisms of BMAA in cyanobacteria, eukaryotic phytoplankton and cycads have not yet been disclosed until now.

The BMAA was first identified from cycads (Li et al. 2022) and a simple two-step pathway was hypothesized to produce BMAA from the *β*-substituted alanine under the action of cysteine synthase-like enzyme and methyltransferase (Brenner et al. 2003). The BMAA molecular structure was also found in an antibiotic component, galantin I, isolated from *Bacillus* (Sakai and Ohfune 1990), in which the 2,3-diaminopropanoic acid (2,3-DAP) was hypothesized as the precursor that was methylated to form BMAA under the catalysis of methyltransferase (Nunn and Codd 2017). In addition, some key cellular processes were suggested to be responsible for biosynthesis of BMAA-containing proteins in diatom cells, including the inhibition of ubiquitination and autophagy function, the decreased accuracy and efficiency of COPⅡ vesicle transport (Li et al. 2022). It should be noted that the BMAA detected in laboratory cultures of diatoms was exclusively in the form of protein-bound precipitates (Wang et al. 2021; Li et al. 2022), whereas BMAA was mainly in free form in cyanobacteria (Yan et al. 2020). The discrepancy between the BMAA forms in diatoms and cyanobacteria may indicate that the BMAA biosynthetic pathway in diatoms was different from that in cyanobacteria (Li et al. 2022).

Harmful algal blooms formed by diatoms showed that the supply of iron (Fe) played a critical role in the development of extensive diatom blooms in coastal upwelling regimes (Bruland et al. 2001). In addition, Fe, as a micronutrient required for phytoplankton growth, contributed to the marine primary productivity and carbon export (Vivado et al. 2022). However, the concentration of Fe in seawater, especially in surface seawater was very low (Hogle et al. 2018), and Fe deficiency in seawater limited the photosynthesis and growth of phytoplankton, resulting in insufficient utilization of nitrogen (Bruland et al. 2001). Furthermore, during periods of low iron availability, the expression and activity of iron-dependent nitrate and nitrite reductases decreased, contributing to a reduction in nitrogen assimilation (Maniscalco et al. 2022).

Iron deficiency may lead to ocean acidification, by the increase of CO_2_ and the decrease of pH in seawater. Therefore, exploring the changes of diatom toxin production under the condition of iron deficiency is essential to discover the biosynthesis mechanism of BMAA and maintain the balance of marine ecosystem. In the present study, the production of BMAA-containing proteins was significantly stimulated and was universal when diatom growth under Fe deficiency stress. RNA-seq data were further used to analyze and explore the related genes and regulatory pathways of BMAA biosynthesis in diatom cells under Fe limitation. The subcellular distribution of BMAA-containing proteins in diatoms was also discussed here. Finally, we suggested the possible reaction process from cysteine and methylamine to form BMAA *in situ*.

## Results and Discussion

### Stimulation of BMAA biosynthesis by Fe limitation in diatoms

Growth of four diatom species was significantly inhibited by Fe limitation (1/3 × Fe) (Fig. S1), however, the corresponding BMAA yield was increased by nutrient stress, maximum up to 7.7 fold in the culture of *T. minima* (Fig. S1). Densities of *T. minima* significantly (p< 0.001) decreased with the decrease of iron concentrations (3 × Fe, 1 × Fe, 1/3 × Fe, 1/6 × Fe) spiked in the culture medium, starting from day 4 onwards (Fig. 1A). This phenomenon is consistent with the previous studies that the supply of Fe stimulated phytoplankton growth and photosynthesis (Thomas 2003; Lelong et al. 2013). The availability of Fe by diatoms was verified by the significant decrease of Fe concentration in culture media during the batch culture period (Fig. 1B), as well as the discrepancy of Fe concentrations in diatom cells cultured with different Fe concentrations (Fig. 1C). Cell size was significantly increased, but Chl *a* content was obviously decreased in diatoms cultured with Fe restriction (Figs. 1D, E). We speculate that the diatom *T. minima* showed a response of cell plasticity to Fe deficiency to adsorb more Fe ions and light capture via increasing cell surface. Notably, BMAA production in *T. minima* was largely improved up to 7.7 fold higher than that of control group by the limitation of Fe (1/3 × Fe) (Fig. 1F). No significant change occurred in the production of BMAA per cell when the Fe concentration was up-regulated (3 × Fe).

**Figure 1.**
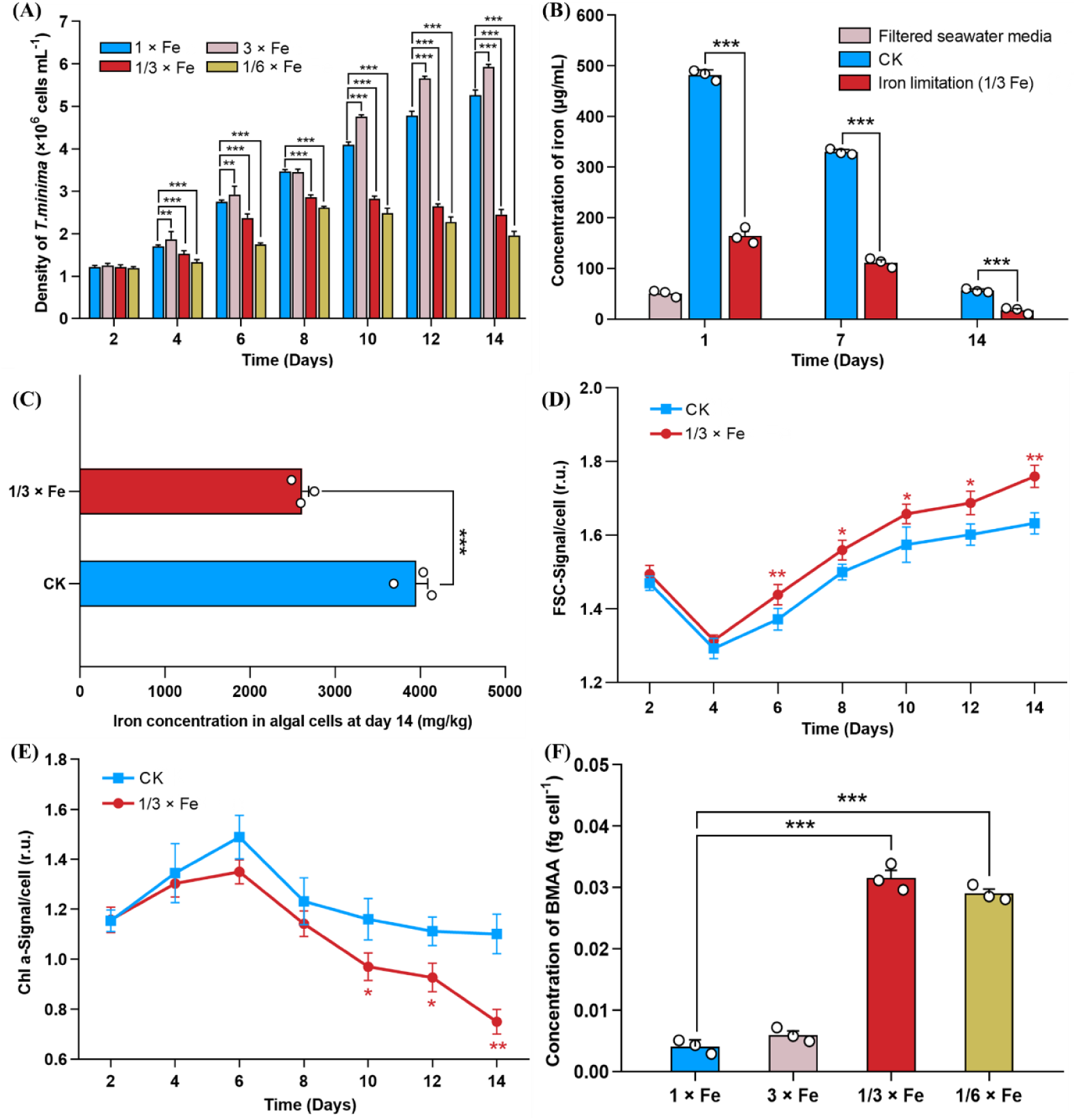
Growth status, determination of the iron concentration and the β-N-methylamino-L-alanine (BMAA) index of the diatoms (*T. minima*) under iron limitation. (A) Average densities of the diatoms in the f/2 medium spiked with different iron concentrations (n = 3): the proportion of iron-containing nutrients decreased by 3-fold (1/3 × Fe), 6-fold (1/6 × Fe) and increased by 3-fold (3 × Fe) and a normal culture has been carried out on f/2 medium (1 × Fe, CK). (B) Changes in the iron concentrations of diatoms were measured in 400 mL of microalgal suspension (n=3). (C) Microalgal cells were collected for iron concentration on day 14 (n=3). (D) Mean Forward Scattered (FSC) signals of diatoms in iron limitation (1/3 × Fe) and CK groups (n = 3). (E) Chlorophyll a fluorescence signal of diatoms in iron limitation group (1/3 × Fe) and CK group (1 × Fe) (n = 3). (F) Changes in the BMAA concentrations of diatoms in the CK group (1 × Fe) and with a different ratio of Fe concentration groups (n=3). The asterisk denotes a statistical difference between the iron limitation group and the control group (one-way ANOVA or Welch’s t-test, * 0.01 < p < 0.05, ** 0.001 < p < 0.01, ***p < 0.001).

In our previous study, BMAA production was also significantly increased in *T. minima* co-cultured with cyanobacteria (Li et al. 2022). The Fe deficiency possibly occurred for diatoms in the co-culture system because cyanobacteria have an extremely high demand for Fe during growth (Morrissey and Bowler 2012; Yong et al. 2022). For another phycotoxin domoic acid (DA), Fe deficiency was thought to enhance DA production of *Pseudo-nitzschia* diatoms and release from cells because DA could resemble a phytosiderophore to enhance the uptake and utilization of Fe (Rue and Bruland 2001; Lelong et al. 2012). Reportedly, the decrease in cell size of *Pseudo-nitzschia* spp. over time may lead to a decrease in the amount of DA per cell (Bates 2018). We hypothesized that the increase in cell size of *T. minima* caused by Fe limitation would lead to an increase in BMAA-containing proteins accumulation per cell in the present study. The biosynthesis of BMAA-containing proteins is possibly an adaptive response to Fe limitation in environment for diatoms like the function of DA.

### Total response of diatom to Fe limitation at transcriptional expression

Raw transcriptome data can be downloaded from the National Center for Biotechnology Information (NCBI) website, PRJNA946367. Quality parameters of the filtered *T. minima* clean reads were shown in Table S1. Iron limitation caused a significant response at the transcriptional level in *T. minima* (Fig. S2). A total of 26,476 differentially expressed genes (DEGs) occurred between experimental group and control group (Fig. 2A), in which 12,279 and 14,197 genes were up- and down-regulated (|log_2_fold change| ≥2), respectively. Kyoto Encyclopedia of Genes and Genomes (KEGG) database was used to enrich all DEGs and the top 20 KEGG enrichment pathways of significant DEGs (|fold change| > 2) were shown in Fig. 2C. The top 20 Genetic ontology (GO) pathways (Padjust < 0.05) were mainly enriched in biological process, cellular component and molecular function (Fig. S3).

**Figure 2.**
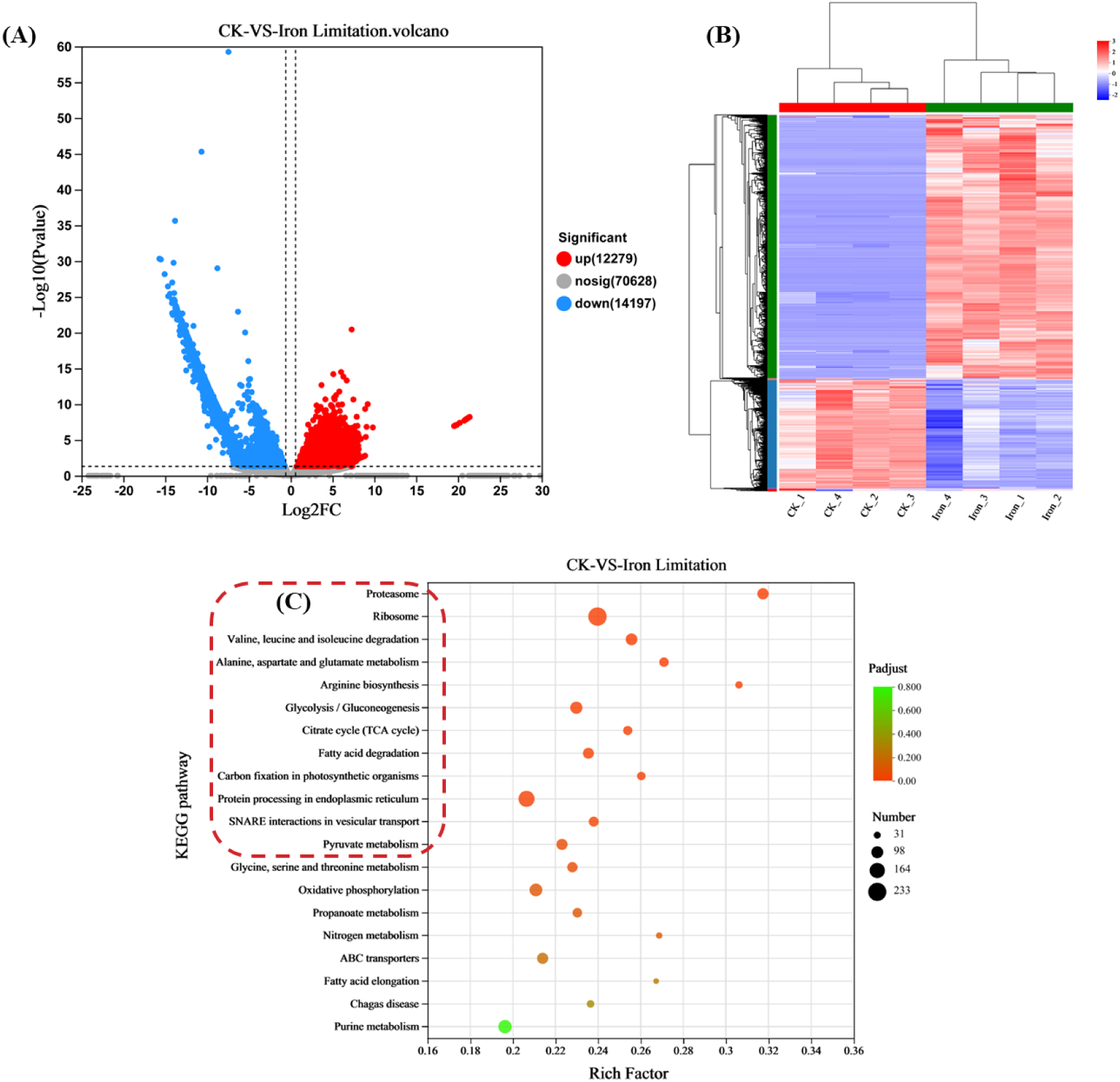
Results of transcriptional analysis of diatoms cultured with normal f/2 condition (CK groups) and iron limitation groups (1/3 ×Fe) in four duplicates. (A) Volcano map of differentially expressed genes (DEGs) in the iron limitation groups (1/3 × Fe) and CK groups (1 × Fe). (B) Cluster analysis of DEGs in eight diatoms samples in the iron limitation groups (1/3 × Fe, Iron_1 to Iron_4) and CK groups (1 × Fe, CK_1 to CK_4). (C) Scatterplot of the top 20 KEGG annotations with P-adjust for the DEGs in diatoms from iron limitation groups (1/3 × Fe) vs. CK groups (1 × Fe).

There were twelve KEGG pathways with Padjust < 0.05 in the top 20 pathways enriched by the DEGs (Fig. 2C). These KEGG pathways were mainly related to energy metabolism, protein biosynthesis and carbon fixation functions in diatoms. Diverse amino acid metabolism pathways related to citrate cycle (TCA cycle) were observed, such as Valine, leucine and isoleucine degradation (map00280; Padjust 0.00006), Alanine, aspartate and glutamate metabolism (map00250; Padjust 0.0005), and Arginine biosynthesis (map00220; Padjust 0.0007). Energy metabolism pathways including glycolysis, TCA cycle, and oxidative phosphorylation made responses to the Fe limitation at transcriptional level. Four KEGG pathways related to protein biosynthesis function were enriched with Padjust < 0.05, including Proteasome (map03050; Padjust 1.33e-9), Ribosome (map03010; Padjust 3.49e-9), Protein processing in endoplasmic reticulum (map04141; Padjust 0.013), and SNARE interactions in vesicular transport (map04130; Padjust 0.022). The production of BMAA-containing proteins possibly resulted from these KEGG pathways related to protein biosynthesis functions.

### Response of carbon fixation of diatom to Fe limitation

Only the KEGG pathway Carbon fixation in photosynthetic organisms (map00710; Padjust 0.009) related to photosynthesis functions was enriched (in top 20) in diatoms stressed by Fe limitation in this study. Some key genes involving Calvin cycle and energy metabolism pathways were shown in Fig. 3. Multiple enzymes involved in the Calvin cycle were overexpressed, e.g. Fructose-1,6-bisphosphate aldolase (FBP; K01086), Ribose-5-phosphate isomerase B (rpiB; K01808) and the small subunit of rubisco (rbcS, K01602). While the expression of Phosphoribulokinase (PRK; K00855) was significantly downregulated in the treated diatoms. The enzyme FBP plays an important role in both carbon fixation and glycolytic metabolic pathways, which is closely related to the direction of carbon flow and carbohydrate synthesis. Carbon fixation efficiency would be increased when FBP was overexpressed because oxidative pentose phosphate pathway activity was reduced (Hing et al. 2019). In addition, FBP is a major regulatory site for Glycolysis/Gluconeogenesis because it could be inhibited by AMP and fructose 2,6-diphosphate, but activated by ATP, Citrate, and 3-phosphoglycerate (Grasmann et al. 2019). Ribose-5-phosphate isomerase (RPI) isomerizes ribose-5-phosphate to ribulose-5-phosphate that contributes to the regeneration of the Rubisco substrate (Le Moigne et al. 2020). In addition, RbcS encodes the small subunit of Rubisco and controls the catalytic activity of Rubisco, determining the efficiency of photosynthetic carbon assimilation (Sakoda et al. 2021). PRK plays an important role in the dark response process, and the status of PRK/GAPDH/CP12 complex is associated with rapid and subtle regulation of PRK activity in response to fluctuations in light availability (Howard et al. 2008). Therefore, the up-regulated expression of FBP, rpiB and rbcS enzyme genes in diatoms under Fe restriction would enhance CO_2_ assimilation efficiency, while the down-regulated expression of PRK gene might be beneficial for the conservation of ATP and NADPH produced during the photoreaction process. The dynamic balance between energy supply and diatom growth was maintained by enhancing carbon fixation.

**Figure 3.**
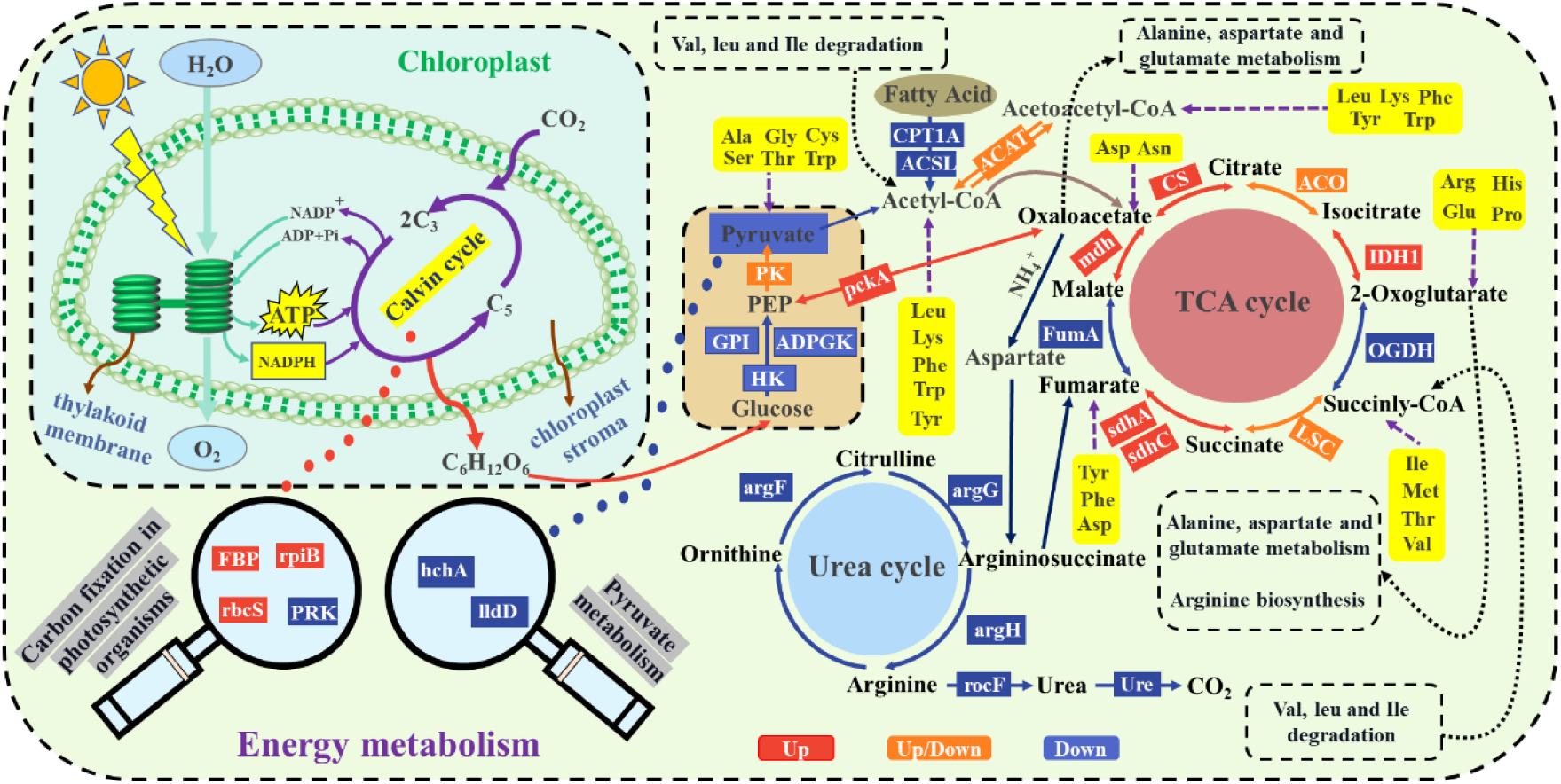
Regulation of energy metabolism-related transcript levels in the diatom *Thalassiosira minima* under iron limitation stress. The red, orange and blue boxes indicate up-regulation, no change and down-regulation of genes, respectively (see text explanation).

### Response of energy metabolism of diatoms to Fe limitation

The whole energy metabolism processes comprised by glycolysis, TCA cycle, and oxidative phosphorylation were markedly affected by the Fe limitation stress in diatom at the transcriptional level. Especially TCA cycle as the core of energy metabolism of aerobic organisms was significantly changed by the stress in diatoms (Fig. 3). The TCA cycle was initiated and mediated by citrate synthase (CS; K01647) to form the first product citrate entering this cycle. Oxaloacetate, as the key reaction substrate catalyzed by CS, can be transformed through malate dehydrogenase (mdh; K00024). The input of acetyl-CoA keeps the TCA cycle circulating to produce ATP, high energy electrons, and release CO_2_ (Sweetlove et al. 2010). However, the supplier of acetyl-CoA from glycolysis to produce pyruvate would be suppressed in diatoms under Fe limitation because some key enzymes involved the glycolysis, such as glucose phosphate isomerase (GPI; K01810), ADP dependent glucokinase (ADPGK; K08074) and hexokinase (HK; K00844), were down-regulated expression in this study (Rengaraj et al. 2012). The downregulation of glutathione-independent glyoxalase (hchA; K05523) and L**-**lactate oxidizing enzymes (lldD; K00101) during pyruvate metabolism also suggested that glycolysis was insufficient for the TCA cycle. Acetyl-CoA for energy metabolism was also supplied by the degradation of fatty acids and ketogenic amino acids such as lysine (Lys) and leucine (Leu) (Sweetlove et al. 2010). Some key genes involved in fatty acid degradation, such as long chain acyl-CoA synthetase (ACSL; K01897) and carnitine palmitoyl transferase 1A (CPT1A; K08765), were also downregulated, indicating that fatty acids were insufficient to maintain the acetyl-CoA balance. In a previous study, evolutionary changes in transcriptional regulation of acetyl-CoA-related pathways, including lipid and branched-chain amino acid metabolism, were used by *Chaetoceros* sp. to balance photosynthetic light capture with changes in metabolism and growth temperature (Liang et al. 2019). In the current study, the expression of CS, mdh, isocitrate dehydrogenase 1 (IDH1; K00031), and succinate dehydrogenase genes (sdhA; K00239, and sdhC; K00241) were significantly up-regulated, but 2-oxoglutarate dehydrogenase (OGDH; K00164) and fumarase A (fumA; K01676) were down-regulated. Therefore, the supply of TCA cycle intermediates may exhibit an off-cycle flux pattern to compensate for carbon losses in the cycle, including other biosynthetic processes that provide a carbon skeleton (Sweetlove et al. 2010). The downregulation of OGDH and fumA enzymes in the treated diatom may be coordinated with the increased photosynthesis (Sweetlove et al. 2010).

To compensate the insufficient acetyl-CoA, the amino acid degradation and synthesis processes were also activated in the diatom in this study. The degradation of valine, leucine, and isoleucine could produce acetyl-CoA, succinyl-CoA, acetoacetyl-CoA, propanoyl-CoA and methyl-malonyl-CoA. The valine, leucine and isoleucine degradation (map00280; Padjust 0.00006) pathway was also enriched in treated diatoms in our previous study (Li et al. 2022). Most genes involved in this amino acid metabolism pathway were significantly up-regulated at transcriptional level, such as 3-hydroxy-3-methylglutaryl-CoA lyase (HMGCL; K01640), recombinant acetoacetyl coenzyme A synthetase (AACS; K01907), alanine glyoxylate aminotransferase 2 (AGXT2; K00827) and methylmalonyl-CoA epimerase (MCEE; K05606). Meanwhile, several key synthetases associated with arginine biosynthesis were down-regulated, including argininosuccinate synthase (argG; K01940), argininosuccinase (argH; K01755) and ornithine carbamoyltransferase (argF; K00611). These key cytoplasmic urea cycle genes would directly or indirectly reduce the biosynthesis of arginine (Li et al. 2022). However, most key genes involved in the alanine, aspartate and glutamate metabolism pathway were significantly up-regulated in the treated diatom. For example, the up-regulation of alanine dehydrogenase (ald; K00259) would accelerate the metabolism of pyruvate to L-valine and then promote D-amino acid metabolism. These amino acid metabolism pathways affected by Fe limitation resulted in the significant decrease of amino acid contents in the diatom in this study (Table S2 and Fig. S4B). The ATP content tested in diatom cells showed that the function of TCA cycle was overall improved to produce more ATP molecules in diatoms under Fe limitation (Fig. S4A).

### Response of protein biosynthesis in diatoms to Fe limitation

Ribosome plays an important role to decode genetic information and produce peptides in endoplasmic reticulum, in which the small subunit of ribosome is responsible for recognizing and decoding the genetic information of mRNA. Most genes involved in the small subunit of ribosome were up-regulated (Fig. 4), which would increase the Rubisco’s catalytic efficiency and specificity (Wang et al. 2001; Spreitzer 2003). However, some genes associated with large subunit of ribosome were significantly down-regulated, suggesting that the transport capacity of target amino acids to form polypeptide chains was reduced (Cong and Shuman 1992). The polypeptide chains formed by ribosome will be transferred from the endoplasmic reticulum (ER) to the Golgi apparatus for further modification. The mannosyl-oligosaccharide glucosidase (MOGS; K01228) was up-regulated in the ER, which involved the shear modification of N-glycans (Sadat et al. 2014). Most proteins will be modified by sugar or sugar chain after translation, and mannose trimming of the glycans triggers the ER-associated protein degradation pathway (Kikuma et al. 2022). The generated monosaccharide glycosides (G1M9) (Fig. 4), as the important folding intermediates, were recognized by ER lectin chaperones calnexin or calreticulin, which cooperate with ER oxidoreductase (ERP57; K08056) to promote protein folding (Kikuma et al. 2022). In this study, the down-regulated expression of ERP57 suggested that the correct folding efficiency of proteins might be reduced, and more misfolded proteins may accumulate. Subsequently, glycoproteins derived from the M9 sugar type were transferred to the Golgi apparatus for further processing (Kikuma et al. 2022), and LMAN1 (K10080) overexpression increased cargo protein transport from ER to ER-Golgi intermediate compartment (ERGIC) and Golgi bodies (Hao et al. 2014).

**Figure 4.**
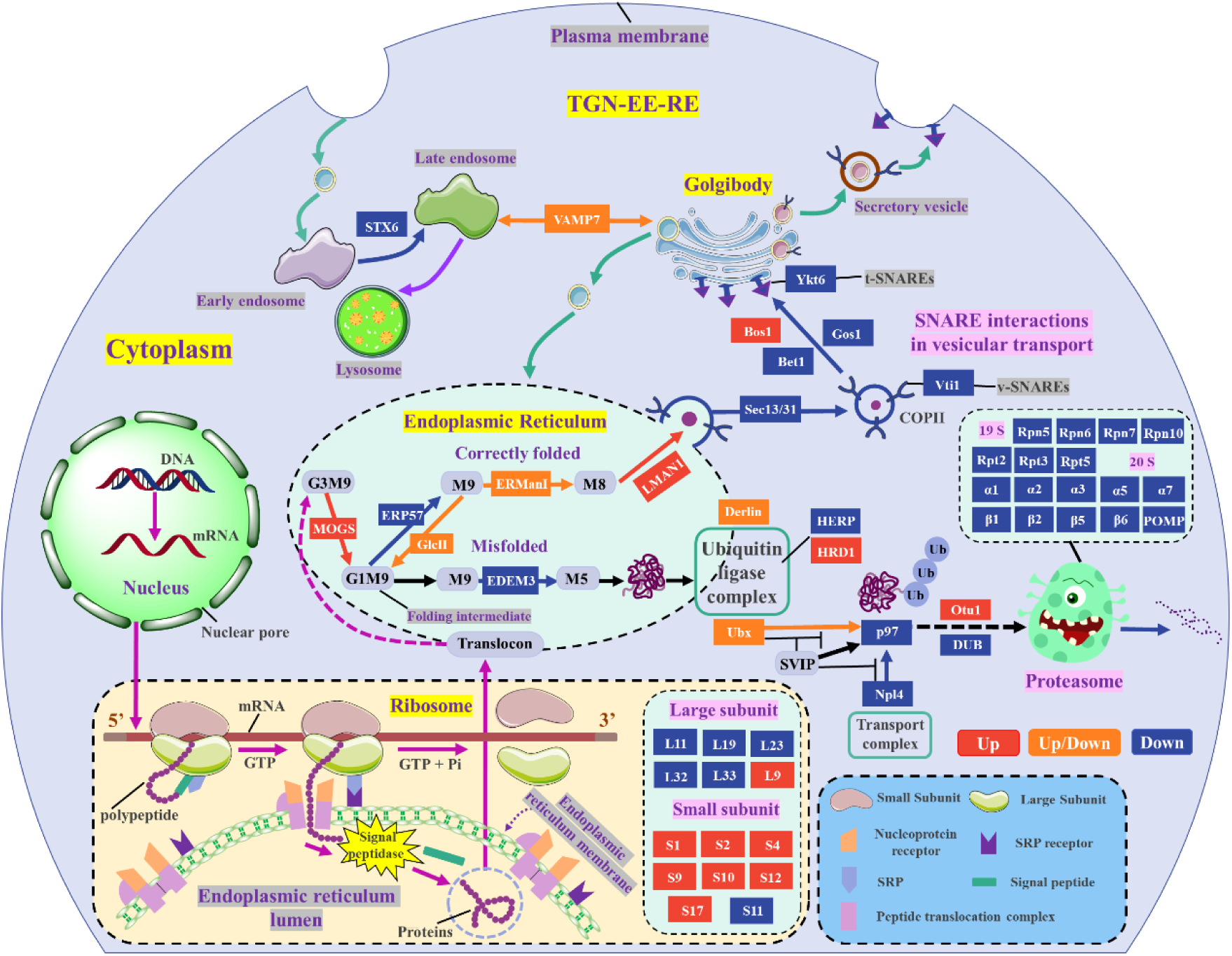
Regulation of protein biosynthesis-related transcript levels in the diatom *Thalassiosira minima* under iron limitation stress. The red, orange and blue boxes indicate up-regulation, no change and down-regulation of genes, respectively (see text explanation). TGN = trans-Golgi network; EE = early endosome; RE = recycling endosome.

Specific uptake of secretory and vesicle targeting (V-SNARE) proteins into coat protein complex II (COPII)-coated vesicles play important roles in the protein transport from ER to Golgi apparatus (Li et al. 2022). The cytoplasmic components of the COP II coat include small GTPase Sar1 and two heterodimeric protein complexes Sec23/24 and Sec13/31 (Barlowe 2002). Activated Sar1-GTP first binds to the membranes and recruits the Sec23/24 complex and then the Sec13/31 complex, inducing polymerization of the coat and deformation of the membranes into buds and vesicles (Barlowe 2002). In this study, the down-regulation of Sec13/31(K14004 and K14005) would lead to a decrease in the activity of the Sar1p GTPase, mediated by Sec23 (Barlowe 2002). COPII vesicles then interact with a select set of COPII vesicle–associated SNAREs (soluble N-ethylmaleimide–sensitive factor attachment protein receptors) to form a *cis*-SNARE complex that facilitates targeting to the Golgi apparatus (Allan et al. 2000). However, in the present study, the transcriptional expression of both genes encoding *t*-SNARE (Vti1; K08493) and *v*-SNARE (Ykt6; K08516) was down-regulated in the treated diatom, indicating a decrease in the accuracy and efficiency of COPII vesicles in the transport of specific proteins (Li et al. 2022). Besides, the efficiency of ER-Golgi transport may be positively influenced by overexpression of BOS1 (K08496), whereas downregulation of BET1 (K08504) and GOS1 (K08495) reduces the capacity of vesicles to be transported from the ER to the Golgi complex (Newman et al. 1992). In addition, Syntaxin6 (STX6; K08498), which was involved in intracellular vesicle trafficking, particularly in the trans-Golgi network (TGN)-early endosome (EE) and recycling endosome (RE) systems, was down-regulated, which may reduce the efficiency of autophagy in removing unwanted molecules such as misfolded proteins (Li et al. 2022). Based on the regulation of the above genes at transcriptional level, the accuracy of transporting proteins from the ER to the Golgi apparatus was weakened, which would lead to misfolded protein accumulation.

All alive cells are burdened with the task of getting rid of misfolded or damaged proteins (Ibarra et al. 2021). Glc residues reattach to the glycan and regain their binding affinity to calnexin and calreticulin if folding was incomplete. Simultaneously, the M9 glycoform was converted to M5 by the transport and trimming of EDEM3 (K10086) (Kikuma et al. 2022), and EDEM3 was responsible for the recognition of misfolded proteins (Polla et al. 2021), the down-regulation of EDEM3 will easily cause the accumulation of incomplete or misfolded glycoproteins in the endoplasmic reticulum (Manica et al. 2021). Terminally the misfolded polypeptides in the ER were retro-translocated to the cytosol, where they were eventually degraded by the ubiquitin-proteasome (UPP) pathway, a process termed ER-associated degradation (ERAD) (Fujimori et al. 2013). In this process, several ER-localized E3 ubiquitin ligases target ERAD substrate proteins for ubiquitination, forming the ubiquitin ligase complex. Where the N-terminal of homocysteine-induced endoplasmic reticulum protein (Herp; K14027) acts as a molecular chaperone to inhibit protein aggregation, while its C-terminal functions as an E3 ubiquitin ligase to promote the degradation of misfolded proteins through the UPP (Luo et al. 2018). As shown in Fig. 4, HERP down-regulation may lead to misfolded protein aggregation and weakened UPP pathway. Fortunately, HERP was able to rapidly stimulate HRD1-mediated ubiquitylation and degradation of aberrant ER proteins(Kny et al. 2011), so overexpression of (HRD1; K10601) will positively respond to the ubiquitination and degradation of misfolded proteins (Kaneko et al. 2012). P97 (VCP; K13525) was an ATPase domain-containing protein cleavage enzyme essential for post-ubiquitylation events in the ubiquitin-proteasome pathway, requiring rapid engagement of ubiquitin ligase-chaperone pairs that restore the ubiquitin signal for proteasome targeting (Hu et al. 2020), and then upon binding to a polyubiquitinated substrate, Npl4 (K14015) in complex with p97 can unfold the ubiquitin to initiate the processing of the substrate (Hu et al. 2020). In our study, most of the genes involved in the transport complex were downregulated, indicating a weakened transport capacity of ubiquitylated proteins. Deubiquitinase (DUB; K11863) and ovarian tumor domain-containing protein 1 (Otu1; K13719) can prevent unwanted ubiquitination to enhance substrate specificity for an associated ubiquitin ligase partner and to promote ER quality control (Zang et al. 2020). Thus, the up-regulation of Otu1 and down-regulation of DUB can effectively regulate the necessary protein for ubiquitination labelling to be efficiently degraded by proteasomes. Ubiquitination was known as an indicator of protein degradation by the 26S proteasome (Li et al. 2022), which comprises two multi-subunit subcomplexes as follows: 20S core particle and two 19S regulatory particles. The 20S was the core of all proteasome conformations and was the major protease responsible for degrading oxidative proteins, while the 19S can degrade ubiquitinated proteins through ATP-dependent mechanisms (Raynes et al. 2016). In addition, POMP (K11599) facilitates the key steps in the formation of the 20S core complex at the ER, coordinating the assembly process and delivering freshly formed proteasomes to the cells at their site of function (Fricke et al. 2007). As shown in Fig. 4, the expression of the α-ring and β-component subunits in the 20S core particle of the proteasome was down-regulated, and the gene of POMP associated with the formation of the 20S core complex was also down-regulated. Furthermore, it cannot be ignored that the expression of genes related to 19S was also down-regulated. Therefore, the role of the UPP in ubiquitin-mediated protein hydrolysis was weakened by inhibiting of proteasomes in diatom under Fe limitation, which could reduce autophagy of abnormal proteins such as BMAA-containing proteins, as demonstrated in our previous study (Li et al. 2022).

### Subcellular distribution of BMAA-containing proteins in diatoms

To locate the BMAA-containing proteins in diatom cells, different organelles were obtained through differential centrifugation and analyzed for BMAA content. Results showed that the BMAA-containing proteins were exclusively detected in the Golgi apparatus and endoplasmic reticulum (Fig. S5). Content of BMAA-containing proteins in *T. andamanica* was approximately 0.025 fg cell^-1^ when the diatom grew to the stable growth stage. The distribution ratio of the BMAA in the individual organelles was evaluated after a differential centrifugation of the homogenate. Content of BMAA in the intact cells was about 0.025 fg cell^-1^ (×150 g, 10 min), and the content of BMAA in ER and Golgi apparatus was also about 0.025 fg cell^-1^ (×100000 g, 90 min). To determine whether the absence of BMAA in other organelles might be due to nuclear or mitochondrial membrane damage, and the presence of BMAA in the Golgi apparatus and endoplasmic reticulum might be due to contributions from other damaged organelles, the integrity of organelle membranes obtained by differential centrifugation were checked by TEM. Membranes of the nucleus, mitochondria, Golgi apparatus, and endoplasmic reticulum were almost complete (Fig. S6), which demonstrated that the BMAA-containing proteins were formed and accumulated in the Golgi apparatus and the endoplasmic reticulum.

### Speculative biosynthesis mechanism for BMAA-containing proteins in diatoms

As described above, the BMAA-containing proteins accumulation was stimulated in diatom by Fe limitation, which is like the change in production of DA toxin by *Pseudo-nitzschia* species under Fe deficiency stress (Lelong et al. 2012). The discovery of the biosynthetic pathway and the gene cluster encoding the phycotoxin DA in *Pseudo-nitzschia multiseries* provided an important methodology for this study to explore the genes involved in the biosynthesis of BMAA (Brunson et al. 2018). Combining our previous study results (Li et al. 2022) and the current study data, two identical KEGG pathways were enriched with low Q values in both studies, including valine, leucine and isoleucine degradation and SNARE interactions in vesicular transport. In addition, four KEGG pathways enriched in the current study were also highly correlated with that reported in the previous study, namely arginine biosynthesis; alanine, aspartate and glutamate metabolism; protein processing in endoplasmic reticulum; and proteasome. For the first time, it was verified that the BMAA-containing proteins were formed and accumulated in the endoplasmic reticulum and the Golgi apparatus of diatom cells. The reduced synthesis of amino acids should contribute to the biosynthesis of BMAA-containing proteins to some degree.

No free form of BMAA was detected in the diatom cells cultured in laboratory. We speculated that the BMAA structure should be changed from other amino acids that have been inserted in some peptides and thus formed the misassembled proteins. Monomethylamine (MMA), dimethylamine (DMA) and trimethylamine (TMA) were relatively common low-molecular weight organic nitrogen compounds in the marine environment (Yang et al. 1994). Marine phytoplankton, such as diatoms, have been shown to take up and accumulate MMA (Gibb et al. 1999). Notably, methylamine in seawater is not only a product of degradation, excretion and metabolism by marine animals and bacteria (Yang et al. 1994), but interestingly, it can also be produced by diatoms (Gibb et al. 1999). In this study, it was found that when *T. minima* growth was stressed by methylamine, *T. minima* density was repressed as growth entered the stationary phase (Fig. S7A), but BMAA synthesis was stimulated (Fig. S7B). When the concentration of methylamine was 0.12 mM, BMAA production increased 2.1-fold (Fig. S7B). Similarly, other studies have found that the addition of Tris, a cell membrane permeable primary amine, had a significant effect on DA production by *Nitzschia pungens f. multiseries* (Douglas DJ et al. 1993), speculating that Tris may be used as a nitrogen source to promote DA production through modification of the enzyme response (Douglas DJ et al. 1993).

CysK serves as a key cysteine synthase, acting as a nucleophilic reagent that can displace sulphide in the β-substitution reaction of amino acids, resulting in the formation of unnatural β-substituted L-α-amino acids (Maier 2003). Five selected genes were compared between the transcriptome sequencing and qPCR gene expression changes (Fig. S4C). The key target gene cysK was upregulated in the current study and our previous study (Li et al. 2022). Interestingly, in several of our selected BMAA-producing diatoms, a positive correlation was also found between BMAA production and cysK gene expression levels (Table S3 and Fig. S7C). Combined these results suggested that the amino acid cysteine inserted in peptide chains could serve as a substrate for nucleophilic reactions with methylamine catalyzed by the cysK enzyme, thus resulting in the production of BMAA-containing proteins (Fig. 5). Furthermore, the decreased ubiquitination-mediated proteolysis and the decreased vesicle trafficking precision further aggravated the accumulation of BMAA-containing proteins in diatom cells. This is the first time to explain the biosynthesis mechanism for BMAA detected in marine diatoms. Meanwhile, this study also hints that the production of BMAA-containing proteins should be increased due to the prevalence stress of Fe limitation in the worldwide coastal waters in the future.

**Figure 5.**
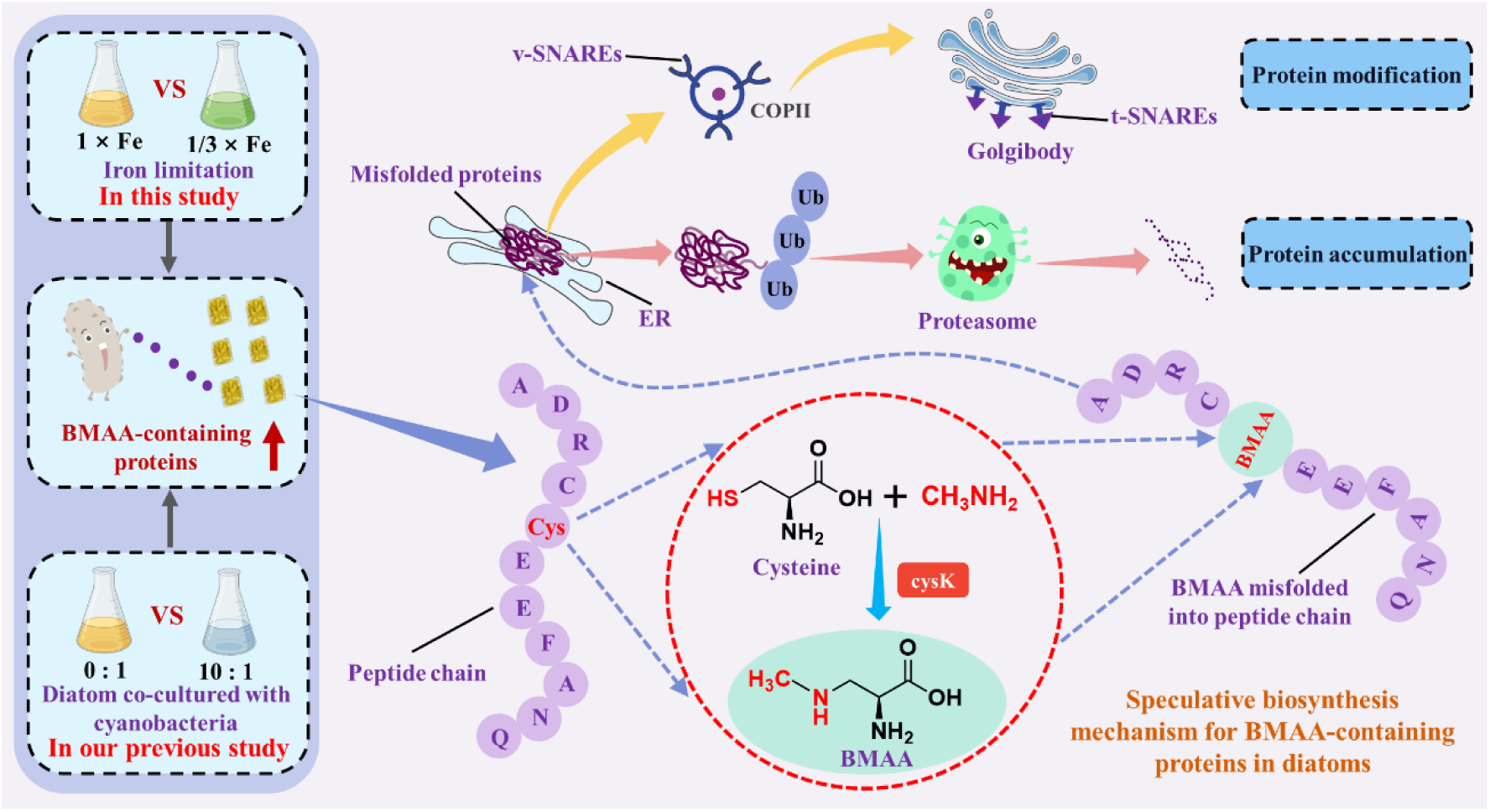
The speculative biosynthesis pathways of BMAA-containing proteins in diatoms.

## Materials and Methods

### Culture of diatoms and iron limitation treatment

Four different diatom species were used in this experiment. One diatom species was isolated from the coast of the East China Sea: *Thalassiosira andamanica* (MC6523), coast of Taiwan Strait; three diatom species were isolated from the coast of the South China Sea: *T. allenii* (MC6441), coast of Hong Kong, *T. gravida* (MC6463), and *T. minima* (MC6475), from the coast of Shantou. All diatoms were cultured in laboratory using Guillard’s f/2 medium. The illumination was LED light at 6000 lux with a cycle 12 h light: 12 h dark, and by gently swirling the 500-mL culture flasks three times daily.

These diatom strains were cultured under iron restriction. The initial density of diatoms was 1.0 × 10^4^ cells mL^-1^. The growth curve was measured to evaluate the effect of iron restriction on their growth, then the diatom cells were collected for BMAA analysis during the stationary growth phase.

The most sensitive diatom *T. minima* to iron limitation was selected to further investigate the effect of different concentrations of iron on the production of BMAA. The initial density of *T. minima* was 1.0 × 10^4^ cells mL^-1^, and four experimental groups were designed corresponding to four different nutrient conditions: the proportion of iron-containing nutrients decreased by 3-fold (1/3 × Fe), 6-fold (1/6 × Fe) and increased by 3-fold (3 × Fe), respectively. In addition, as a control group, the iron content of the f/2 medium was not changed (1 × Fe). All cultures were cultured in 500-mL Erlenmeyer flasks containing 400-mL medium, the cultures were subsampled (1 mL) on days 0, 2, 4, 6, 8, 10, 12, 14 to calculate their growth rates, and were harvested on day 14 for BMAA analysis. Moreover, Fe concentration, ATP content, amino acid content, and transcriptome were analyzed under 1/3 × Fe cultivation of *T. minima*.

Flow cytometry was used to assess the cell density, morphological and physiological measures in all cultures (Li et al. 2022). Specifically, diatom cells were counted and observed in all experiments using an Accuri C6 plus flow cytometer (BD Biosciences, NJ, USA). Fluorescence of chlorophyll a (Chl a) was used to measure the density of at least 50-μL diatom samples, and with an injection flow of 66 μL/min. In addition, cell size and Chl a fluorescence intensity can be characterized by cell forward scattering (FSC) signals.

### Preparation of samples and analytical methods

Extraction of the total soluble form of BMAA: The microalgal cells were centrifuged at 6577 × g for 10 min at 4°C and collected in a 10-mL centrifuge tube when the microalgal cells grew to the stationary growth phase. The collected pellets were stored at -80°C for 24 h and freeze dried in a vacuum freeze dryer (-50°C, 20 Pa), then the centrifuge tube containing the diatom pellets was filled with 5-mL of 0.1 M trichloroacetic acid (TCA). The suspension was frozen in liquid nitrogen for 10 min, thawed at room temperature and this treatment was repeated three times. Diatom cells were then disrupted by sonicating on ice-water mixture for 10 min. The supernatant (2-mL) was transferred to 4-mL glass vials following centrifugation at 6577 × g for 10 min at 4°C, and evaporated under N_2_ gas at 55°C. The residue was dissolved in 1-mL of 6 M HCl and heated at 110°C for 24 h to hydrolyze proteins. After cooling to room temperature, the residue was evaporated again with N_2_, and reconstituted with 1-mL of 20 mM HCl. The extract was purified using Oasis-MCX solid phase extraction (SPE) cartridges (3 cc, 60 mg, Waters, Milford, USA) according to previously described procedures (Wang et al. 2021; Li et al. 2022).

Extraction of the protein-bound form of BMAA: The extraction procedure of protein-bound BMAA was mainly based on the previous study of our group (Wang et al. 2021; Li et al. 2022). Specifically, after the extraction of the soluble BMAA as described above, the precipitated particles were resuspended in 3-mL of 6 M HCl using TCA, transferred to a 4-mL glass bottle, and hydrolyzed at 110°C for 24 h. The solution was then transferred to a 10-mL tube and centrifuged at 6577 × g for 10 min. The supernatant (2-mL) was dried under N_2_ gas at 55°C before reconstituting with 1-mL of 20 mM HCl. The same SPE technique was performed before analysis.

Analysis of BMAA: Analysis of BMAA was performed using a Thermo Ultimate 3000 HPLC (Thermo Fisher Scientific, Bremen, Germany) coupled to an AB-Sciex Qtrap 4500 mass spectrometer (AB Sciex Pte. Ltd, Singapore) with an electrospray ionization source. A binary mobile phase system provided both water (solvent A) and acetonitrile (solvent B), each solvent contained 50 mM (0.1%) formic acid. The target compounds were separated at 30°C on a SeQuant® zwitterionic hydrophilic interaction liquid chromatography (ZIC-HILIC) column (150 mm × 2.1 mm, 5 µm). The mobile phase gradient elution conditions for HPLC gradient elution were shown in Table S4. A flow rate of 350 µL min^-1^ and an injection volume of 5-µL were used with an electrospray voltage of 5500 V and a source temperature of 350°C. Nitrogen was used for the nebulizer and curtain gases. Multiple reaction monitoring (MRM) mode was used to quantify concentrations of BMAA. With the decluster potential (DP) of 30 V and a collision energy (CE) of 13, 11, 17, 24 and 24 eV, five transitions were used, including *m/z* 119 -> 102, 101, 88, 56 and 44, the target BMAA was quantified using *m/z* 119 -> 88.

Extraction of Amino acids: Add 5-mL of 6 M HCl to the centrifuge tube containing the microalgal pellets, then transfer to a 10-mL digestion tube, add high purity nitrogen, and hydrolyze at 110°C for 22 h. After hydrolysis, cool to room temperature, filter the hydrolysate into a 25-mL volumetric flask with filter paper, wash the hydrolysis tube with ultrapure water for several times, filter it with filter paper, combine and transfer it into a 25-mL volumetric flask, use ultrapure water to volume, and mix well. Take 2-mL of the solution after constant volume into a 4-mL sample bottle, dry it with nitrogen at 55°C, and then add 1-mL of 20 mM hydrochloric acid to dissolve it again, after 0.22 μm water system filter membrane was filtered into 1.5-mL sample injection bottle, and the obtained amino acid extract was stored in a refrigerator at -20°C.

Analysis of Amino acids: A TSKgel Amide-80 HILIC column (250 mm × 2.0 mm, 5 µm, Tosoh Bioscience LLC) was used at column temperature 30°C, and amino acids were separated by a binary mobile phase. Mobile phase A was ultra-pure water containing 2.0 mmol L^−1^ formic acid and 50 mmol L^−1^ ammonium formate, and mobile phase B was acetonitrile. The injection volume was 5-µL, and the flow rate was 300 µL min^−1^. The MRM model was used for amino acid analysis (Table S5). Mass spectrum parameters were set as follows: curtain gas pressure (CUR) 40 psi, electric spray voltage (IS) 5500 V, atomization temperature (TEM) 450°C, atomization gas pressure (GS1) and auxiliary gas pressure (GS2) 55 psi and 55 psi respectively, inlet potential (EP) 10 V, and impact chamber ejection voltage (CXP) 12 V. Gradient elution was shown in Table S6.

The concentrations of Fe element in microalgal cells and suspension were measured by ICP-MS (Thermo Fisher iCAP RQ, USA). The ATP content was determined in accordance with the instructions of the scientific kit (Solarbio, Beijing, China).

### RNA-seq analysis and qRT-PCR verification

The transcriptome analysis was performed by Shanghai Majorbio Bio-pharm Biotechnology Co., Ltd. (Shanghai, China). Total RNA was extracted from *T. minima* using TRIzol® reagent (Invitrogen, Beijing, China) according to the manufacturer’s instructions, and genomic DNA was removed using DNase I (Takara, Beijing, China). RNA concentrations ranged from 416 to 845 ng μL^-1^, and 1 μg of RNA from each sample was used for transcriptome sequencing. A fragment analyzer (Agilent 5300 fragment analyzer, M5311AA, USA) was used to assess the quantity and quality of total RNA. The range of RNA integrity number (RQN) values was 8.2-8.8 and 8.0-10.0 in the monoculture and iron-limited culture groups, respectively. Purified mRNA was isolated from total RNA using a Poly(A) mRNA Magnetic Isolation Module according to the manufacturer’s protocol. Then double-stranded cDNA was synthesized using a SuperScript double-stranded cDNA synthesis kit (Invitrogen, CA, USA) with random hexamer primers (Illumina). Following Illumina’s library construction protocol, the synthesized cDNA was then subjected to end repair, phosphorylation and ’A’ base addition. Libraries were size selected for cDNA target fragments of 300 bp on 2% Low Range Ultra Agarose followed by PCR amplified using Phusion DNA polymerase (NEB) for 15 PCR cycles. After quantified by TBS380, paired-end RNA-seq sequencing library was sequenced with the Illumina NovaSeq 6000 sequencer (2 × 150 bp read length).

Use the software Fastp (https://github.com/OpenGene/fastp) to filter the original sequencing data and obtain high-quality sequencing data (clean data). Trinity (https://github.com/trinityrnaseq/trinityrnaseq/wiki) was a relatively authoritative software currently applicable to the assembly of Illumina short fragment sequence. This software integrates three independent software modules to process and splice many RNA-seq data in turn, namely, Inchworm, Chrysalis, Butterfly. To identify DEGs (differentially expressed genes) between two different groups, the expression level of each gene was calculated according to the transcripts per million reads (TPM) method, RSEM (http://deweylab.github.io/RSEM/) was used to quantify gene abundances. Essentially, differential expression analysis was performed using the DESeq2 (http://bioconductor.org/packages/stats/bioc/DESeq2/), DEGs with |log_2_(foldchange)| ≥ 1 and P-adjust ≤ 0.05 were deemed significantly different expressed genes.

The Goatools (https://github.com/tanghaibao/GOatools) software determined the functional categories of gene ontology (GO) terms associated with DEGs with a significant P-adjust < 0.05. The KEGG (Kyoto Encyclopedia of Genes and Genomes) enrichment analysis was carried out by KOBAS (http://kobas.cbi.pku.edu.cn/home.do). To control and calculate the false positive rate, BH (FDR) method was used for multiple tests.

The qRT-PCR was used to detect the effect of iron limitation on the expression level of five genes in *T. minima*, including argH, EDEM3, Bet1, MOGS and cysK. The reference gene database for candidate genes and internal reference genes *β*-action-F and *β*-actin-R primer pairs was designed using Primer Premier 5.0. The primer sequences of the detected genes were shown in Table S7. Relative gene expression level was calculated by the 2−ΔΔCt method.

### Distribution of BMAA in diatom cell organelles

The *T. andamanica* was selected for differential centrifugation (Beckman Coulter, Inc., CA, USA) to analyze the distribution of BMAA in different organelles. Adaptation and optimization of the differential centrifugation method in accordance with the previous studies (Lavoie et al. 2009b; a). Specifically, when the diatom growth entered the stable growth period, the microalgal cells were collected by centrifugation at 4°C and 6577 × g for 10 min, and 0.25 mol L^-1^ sucrose was used as the isotonic homogenate medium for ultrasonic disruption of microalgal cells (50 W, pulse = 0.2 s/s) to obtain a homogenate of different organelles and their contents, and each group was ultrasonically disrupted on ice-water mixture for 4 min. Prior to sonication, they were frozen 3 times with liquid nitrogen. As shown in Fig. S8 and Table S8, various organelles such as nuclei, mitochondria, Golgi bodies, and endoplasmic reticulum were separated by the first precipitating larger particles in the homogenate at low speed and then precipitating particles floating in the supernatant at high speed. Moreover, the content of protein-bound BMAA in organelles obtained at different rotational speeds was evaluated.

Transmission electron microscopy (TEM; ht7770, Hitachi, Japan) was used to evaluate the integrity of each organelle separated by differential speed, and the organelles were fixed in 2.5% glutaraldehyde for 12 h, washed three times with phosphate buffer, fixed in 1% osmium tetroxide for 2 h, then dehydrated with graded ethanol (30, 50, 70, 80, 90, and 100%) and permeated with tert-butyl alcohol. Finally, the samples were lyophilized, gold plated, and observed by TEM.

### BMAA synthetic validation

The monomethylamine used in these experiments was purchased from Chron Chemicals (Chengdu, China). *T. minima* was selected for the methylamine stress test, the culture conditions and initial inoculum density of *T. minima* were the same as in 2.1 above, and when the diatoms were grown in for two days, methylamine was added at the beginning of the logarithmic phase. Four experimental groups were designed corresponding to four different methylamine concentrations: 0.06, 0.12, 0.24 and 0.48 mmol L^-1^, the control group was not added by methylamine. The microalgal density was periodically observed, and the microalgal cells were collected for testing of BMAA content when diatom growth entered a stable period. In addition to the four BMAA-producing diatoms described above, the BMAA-producing diatoms *Chaetoceros lorenzianus* and the non-BMAA-producing diatom *Thalassiosira lundiana* (from the coast of Shantou) were also selected (1 × Fe). These diatoms were also cultured under the same conditions as described above. This was done to verify that there is a linear relationship between the content of BMAA in the different diatoms and its synthesis of the potentially relevant and important target gene, cysteine synthase (cysk). BMAA levels were determined using LC-MS/MS method and the expression of cysK gene was quantified using the qRT-PCR assay.

### Statistical analysis

Four duplicate samples of diatom cultured with normal condition and Fe limitation (1/3 ×Fe) were used for transcriptional analysis. The other experiments were carried out in triplicate. The results were presented as the mean ± standard deviation. The statistical significance was analyzed using one-way analysis of variance (ANOVA) followed by Tukey’s test. Statistically significant was defined as a p-value less than 0.05. All statistical analyses were performed using IBM SPSS 27.

## CRediT authorship contribution statement

**Xianyao Zheng**: Experimental design, Data acquisition and curation, Methodology, and Writing – original draft; **Aifeng Li**: Conceptualization, Methodology, Funding acquisition, Project administration, Supervision, Writing – review & editing; **Jiangbing Qiu**: Methodology, Project administration, Writing – review & editing; **Guowang Yan:** Data acquisition; **Peng Zhao**: Data acquisition; **Min Li**: Data acquisition; **Fanping Meng**: Methodology, Writing-review & editing.

## Acknowledgments

This research was supported by the National Natural Science Foundation of China (U2106205; 41676093).

